# Complement receptor 3 forms a compact high affinity complex with iC3b

**DOI:** 10.1101/2020.04.15.043133

**Authors:** Rasmus K. Jensen, Goran Bajic, Mehmet Sen, Timothy A. Springer, Thomas Vorup-Jensen, Gregers R. Andersen

## Abstract

Complement receptor 3 (CR3, also known as Mac-1, integrin α_M_β_2_, or CD11b/CD18) is expressed on a subset of myeloid and certain activated lymphoid cells. CR3 is essential for the phagocytosis of complement-opsonized particles such as pathogens and apoptotic or necrotic cells opsonized with the complement fragment iC3b and to a lesser extent C3dg. While the interaction between the iC3b thioester domain and the ligand binding CR3 α_M_ I-domain is structurally and functionally well characterized, the nature of additional CR3-iC3b interactions required for phagocytosis of complement opsonized objects remain obscure. Here we analyzed the interaction between iC3b and the 150 kDa headpiece fragment of the CR3 ectodomain. Surface plasmon resonance experiments demonstrated a 30 nM affinity of CR3 for iC3b compared to 515 nM for the iC3b thioester domain. Small angle x-ray scattering analysis revealed that iC3b adopts an extended but preferred conformation in solution. Upon interaction with CR3, iC3b rearranges to form a compact receptor-ligand complex. Overall, the data suggest that the iC3b-CR3 interaction is of high affinity and relies on minor contacts formed between CR3 and regions outside the iC3b thioester domain. Our results rationalize the more efficient phagocytosis elicited by iC3b than by C3dg and pave the way for development of specific therapeutics for treatment of inflammatory and neurodegenerative diseases that do not interfere with recognition of non-complement CR3 ligands.

## Introduction

The complement system is a central part of vertebrate innate immunity. It connects to other branches of the immune system, including adaptive immunity through its functions especially in stimulation of antibody formation. Complement is a tightly regulated proteolytic cascade, which upon activation leads to cleavage of the 186 kDa complement component 3 (C3) into an anaphylatoxin C3a and the opsonin C3b (Fig. 1A). C3b is deposited on the surface of the complement activator through covalent bond formation when an activator nucleophile reacts with an exposed thioester (TE) present in the TE domain of nascent C3b. Host cells present glycans that attract the fluid phase regulator factor H (FH), and also express complement regulators membrane cofactor protein (MCP/CD46) and CR1/CD35. These regulators bind specifically to C3b and enable its degradation by the protease factor I (FI). As a result, C3b is quickly converted to iC3b (1), which acts as a powerful opsonin, as it is recognized by complement receptor 2 (CR2) as well as the two integrin receptors CR3 (CD11b/CD18 or integrin α_M_β_2_) and CR4 (p150,95, CD11c/CD18 or integrin α_X_β_2_). Whereas C3b has a well-defined conformation, double cleavage by FI of C3b in one of two connections to its CUB domain leads to a flexible attachment of the thioester domain to the C3c moiety (Fig. 1A) (2–4). Hydrogen deuterium exchange suggested that in iC3b, the remnants of the degraded CUB domain become surface exposed and disordered (5).

**Figure 1.**
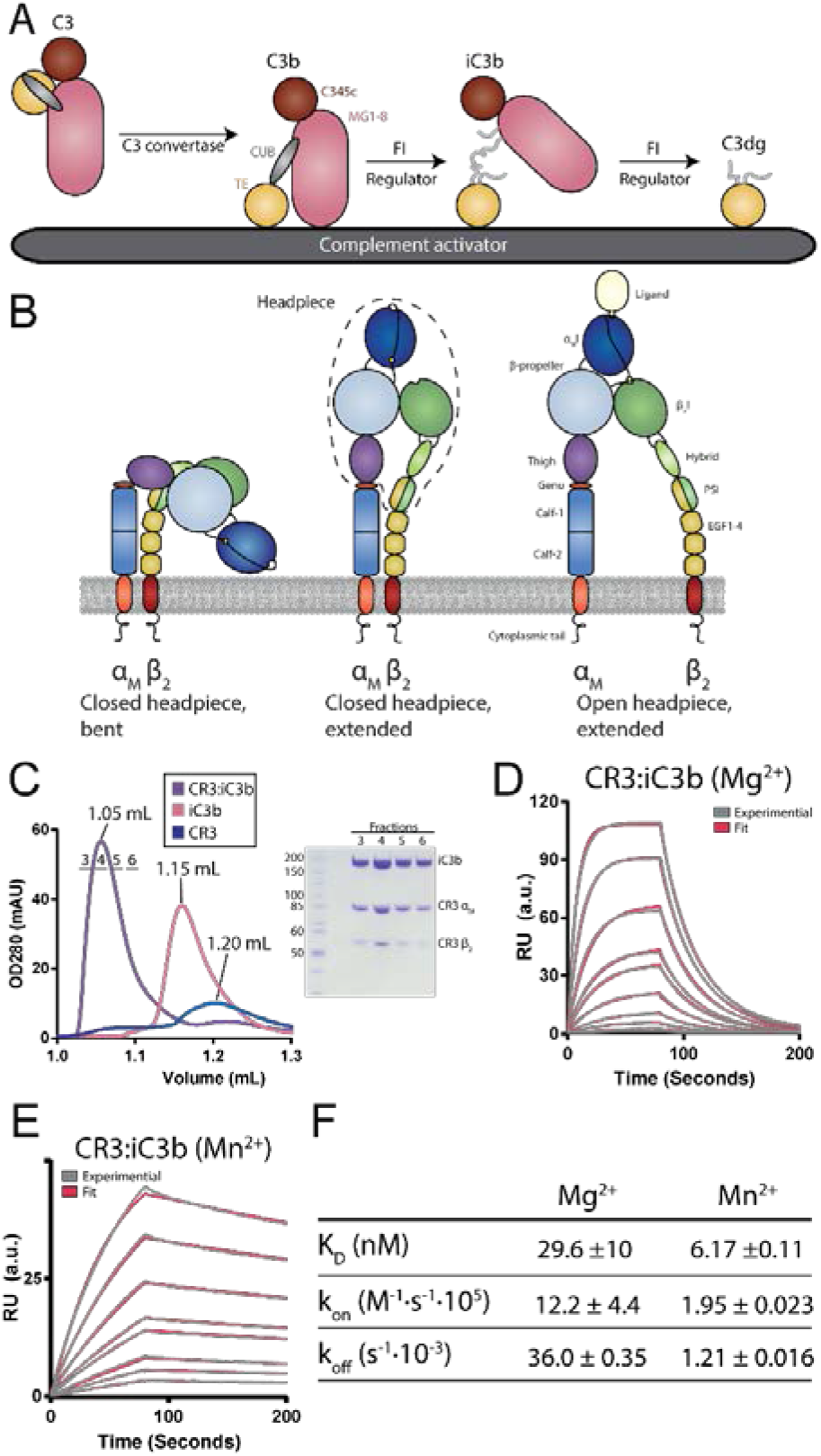
Characterization of the CR3 headpiece fragment. C3 is cleaved by a C3 convertase into the opsonin C3b, which is deposited on the activator surface. C3b can then be further degraded into iC3b by the FI protease, aided by a regulator binding to C3b. FI may further degrade iC3b into C3dg. **B.** CR3 adopts three different overall conformations. The bent conformation (left) and intermediate, extended closed conformation (middle) are low affinity. The extended, open conformation is ~1,000-fold higher affinity (right). The dashed line indicates the headpiece fragment. **C.** SEC analysis of the complex formation between CR3 and iC3b. The complex elutes significantly earlier compared to both iC3b and CR3 alone. The fractions indicated in the chromatogram by a horizontal line is analysed by non-reducing SDS-PAGE. **D.** SPR sensorgrams for the interaction of CR3, injected on an iC3b surface in a Mg^2+^ buffer. CR3 was injected at 100, 50, 25, 12.5, 10, 5, 2.5, 1.25, 0.625, 0.3125 nM. The raw curves are shown in grey and the fit are shown in red. The dissociation constant calculated as K_D_=k_off_/k_on_ is indicated. The on- and off-rates are the average of three independent experiments. **E.** As in panel D, but in a Mn^2+^ buffer, only curves for CR3 concentrations 25, 12.5, 10, 5, 2.5, 1.25, 0.625. 0.3125 nM are displayed. **F.** Table summarizing the kinetic constants determined by SPR, ± the standard deviation.

Recognition of iC3b by CR3 leads to several physiological responses dependent on the cell type and activation state of the CR3-expressing cell, including phagocytosis of dying host cells or pathogens (6, 7). CR3, consisting of the non-covalently associated integrin α_M_ and β_2_ subunits, is highly expressed on the plasma membrane of myeloid cells including macrophages, monocytes, dendritic cells and neutrophil granulocytes and is upregulated from storage granules upon stimulation. Certain lymphoid leukocytes such as natural killer cells and activated T cells also express CR3, and expression is further inducible in other leukocytes (6, 7). CR3 is also highly expressed in microglia, the mononuclear phagocytes of the central nervous system (CNS), where CR3-mediated phagocytosis of iC3b opsonized presynaptic termini of neurons is important for neural development and homeostasis (8–11). *In vivo* studies leave no doubt about the importance of CR3-supported mechanisms, both as a protective agent against infection (12) and as an aggravating factor in diseases with a poorly regulated inflammatory response, for instance, as observed in animal models of multiple sclerosis and Alzheimer’s disease (12, 13). CR3 also plays a key role in complement stimulation of the adaptive immune system. Immune complexes containing complement-opsonized antigens drain with the afferent lymphatics into the subcapsular sinus where complement-opsonized antigens are taken up by subcapsular sinus macrophages via CR3 and carried across the subcapsular sinus floor. Next, the antigen is handed off to non-cognate B cells via complement receptor 2 which transport it into the follicle (14).

CR3, similar to other integrins, adopts at least three distinct conformations in the cell membrane, which control the activity of the protein. These conformations are the bent-closed and extended-closed conformations with low affinity for ligands and the extended-open conformation with high affinity (15) (Fig. 1B). The conformation of integrins is regulated through inside-out signaling, where stimuli received by the cell through other receptors are signaled to the integrin, and outside-in signaling, where a ligand is recognized by the integrinThis signaling, together with the signal of tensile force relayed through the integrin, together stabilize the high affinity conformation (16–18).

In the CR3 α_M_ chain, the I-domain (α_M_I) contains the primary ligand binding site, with a metal ion-dependent binding site (MIDAS) at its center. The I-domain binds a plethora of ligands including iC3b, ICAM-1, RAGE, platelet factor 4, mindin, platelet glycoprotein Ib, sialylated FcγRIIA, CD40L, LL-37, LRP1, fibrinogen, and the LukAB cytotoxin (19–28). Many of these interactions seem at least in part to rely on the ability of the α_M_I to weakly bind glutamate side chains, which translates into avid interactions from multivalent binding of membrane-tethered CR3 (29). By contrast, C3 fragments present distinct high-affinity interactions. We previously established that the major binding site for the CR3 α_M_I is located in the TE domain of iC3b and this interaction is characterized by a dissociation constant (K_D_) of 600 nM (30). However, prior studies suggested that the full-length receptor has a higher affinity for iC3b (31, 32), and other studies (33–36) proposed one or more additional recognition sites between iC3b and CR3. A recent structural analysis by negative stain electron microscopy (nsEM) of the iC3b-CR3 headpiece complex suggested the proximity of regions in iC3b close to the C345c domain and the β-propeller/β I-like domain portion of the CR3 headpiece. However, a three dimensional reconstruction was not obtained and the authors suggested that the tendency of the MG ring with its C345c domain in iC3b and the tendency of the CR3 headpiece to lay flat on EM grids may have broken a 3-dimensional interaction in the vicinity of the domains that were seen to lie close on grids (2).

To investigate whether the iC3b-CR3 complex is an ordered complex with a specific conformation or a flexible ensemble of conformations due to iC3b flexibility, we analyzed the complex between iC3b and the CR3 headpiece in solution through multiple biochemical and biophysical techniques. We now show that the CR3-iC3b interaction is characterized by a 17-fold higher affinity than the minimal complex between the iC3b TE domain and the CR3 αM I-domain (30). SAXS rigid body analysis suggests that the iC3b undergoes a significant conformational change upon CR3 recognition allowing CR3 to interact with regions outside the iC3b thioester domain.

## Results

### The CR3 headpiece forms a high affinity complex with iC3b

The CR3 headpiece was purified from HEK293S GnTI^−^ cells stably transfected with plasmids encoding the α_M_- and β_2_-chains of the 140 kDa CR3 headpiece fragment (dashed outline, Fig. 1B). We investigated the oligomerization state of our CR3 headpiece using analytical SEC in Mn^2+^, Mg^2+^, and Ni^2+^-containing buffers (Fig. S1A). At low concentrations, the CR3 headpiece in both Mg^2+^ and Ni^2+^ mainly eluted as a monomer, but a small dimer fraction was also present. Conversely, in Mn^2+^ CR3 eluted as a dimer in agreement with prior findings that the CR3 headpiece is prone to dimerization in a divalent metal-ion dependent fashion (2). Concentration-dependent dimerization was further confirmed by SAXS (Fig. S1B-E). To verify that the recombinant CR3 headpiece is able to form a stable complex with iC3b, we assessed the stability of the complex by SEC. The CR3:iC3b complex eluted earlier than either of the two individual proteins, in a monodisperse peak (Fig. 1C).

We used surface plasmon resonance (SPR) to measure the affinity and the kinetics of the CR3 headpiece-iC3b interaction. We coupled C3b to biotin through its free thioester cysteine side chain and subsequently converted C3b to iC3b by FI cleavage in the presence of FH as previously described (37). The biotinylated iC3b was then bound to a streptavidin-coated SPR chip surface. The CR3 headpiece bound iC3b in a Mg^2+^ containing buffer with a dissociation constant K_D_ = 30 nM when fitted to a 1:1 interaction model (Fig. 1D+F and Table S1). In a Mn^2+^ containing buffer, CR3 bound to iC3b with a significantly lower k_on_ (Fig. 1E). The dissociation rate was however 30-fold lower than with Mg^2+^, and the CR3 affinity for iC3b in the presence of Mn^2+^ was therefore approximately 5-fold higher with an apparent dissociation constant K_D_ of 6.2 nM (Fig. 1E-F and Table S1).

### Additional iC3b binding sites are present in CR3

The affinity of the CR3 headpiece binding to iC3b was ~18 fold higher than previously described for the isolated CR3 α_M_I domain (30). We therefore investigated whether this was due to stronger binding through the α_M_I domain in the context of the CR3 headpiece or whether CR3 contains one or more additional interaction sites for iC3b outside of the α_M_I domain. For this purpose, we used the recombinant C3d fragment contained within C3dg but with the flexible remnants of the C3g fragment removed (30). As above, the apparent affinity of the CR3-C3d complex was measured using SPR (Fig. 2A-B). Because the binding kinetics were very fast and data could not be robustly fitted to a 1:1 interaction model, we instead performed a steady-state analysis and measured an apparent K_D_ = 515 nM similar to the affinity of the α_M_I domain for iC3b (30). This suggests that embedding of the α_M_I into the CR3 headpiece does not significantly change its affinity for the C3d moiety in iC3b. Next, we performed an SPR-based competition assay where we measured the binding of the CR3 headpiece to immobilized iC3b in the presence of increasing concentrations of free C3b, iC3b and C3d (Fig. 2C-E). Fluid phase iC3b and C3d competed for binding to the immobilized iC3b whereas C3b did not, demonstrating that the competition was ligand-specific. Fluid phase iC3b robustly competed for CR3 binding whereas, by contrast, presence of C3d only produced a marginal decrease in the binding signal even at a 50-fold molar excess of C3d. In summary, our SPR data demonstrated that additional contacts, outside of the α_M_I domain:TE interface described by X-ray crystallography (30), contribute to the CR3 interaction with iC3b.

**Figure 2.**
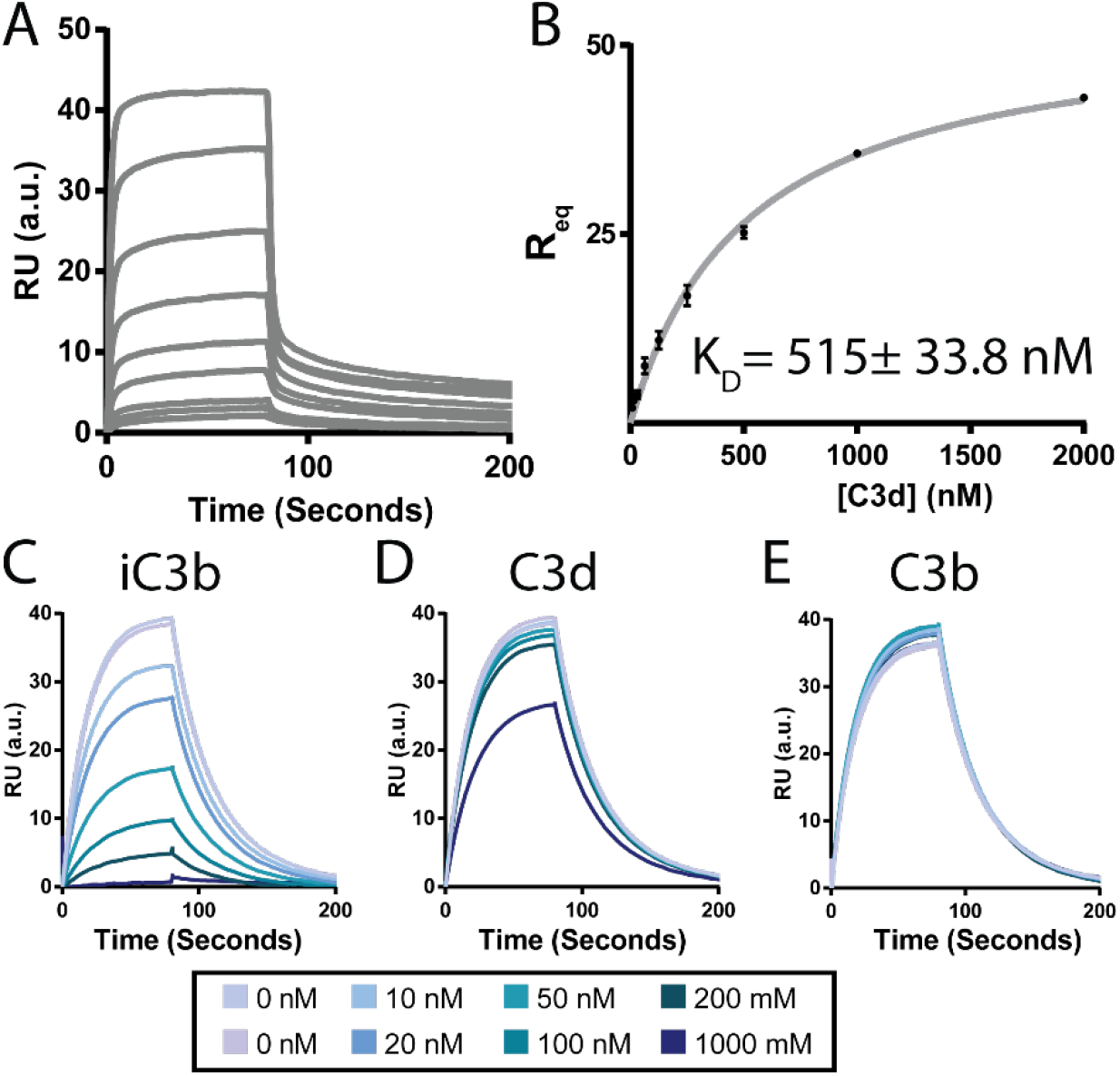
Analysis of the interaction between C3d and CR3. **A.** Sensorgrams from an SPR experiment where the CR3 headpiece at 2000, 1000, 500, 250, 125, 62.5, 31.25, 15.63, 7.81 nM was injected on a C3d coated sensor in a Mg^2+^ buffer. **B.** Steady-state analysis of SPR experiments in panel A. Average values ± the standard deviation for three repetitions are plotted, and the K_D_ value is determined by non-linear regression and given ± the standard deviation. The resulting K_D_ value is 17 fold higher than the K_D_ determined for the CR3:iC3b complex. **C-E.** Sensorgrams of SPR competition assays where 20 nM of CR3 was incubated with variable concentrations of iC3b (**C**), C3d (**D**), or C3b (**E**) before being injected on an iC3b-coated sensor.

### iC3b adopts an extended but stable conformation

After C3b is cleaved by FI to form iC3b, the CUB domain is thought to become disordered and the TE domain no longer closely associates with the MG-ring (2–5). To understand the structural state of the CUB and TE domains after iC3b formation, we recorded SAXS data on iC3b, and for comparison C3b (Fig. S2A). Guinier analysis did not suggest interparticle effects. C3b and iC3b exhibited R_g_ values of 49 Å, and 53 Å respectively (Fig. S2B-C), well in line with earlier reports (3, 38). Comparison of the Kratky plots (Supplementary Fig. S2A) showed that iC3b adopts a more extended structure than C3b. To investigate the solution conformation of iC3b further, we performed rigid body modeling against the scattering data of iC3b. We also performed the same analysis against the data for C3b as a control (Fig. S2D-F). The C3c fragment excluding the C345c domain was modelled as a single rigid body. The C345c domain was modelled independently due to the known flexibility of the domain, but was restrained to maintain the disulphide to MG7. For C3b the CUB and TE domains were treated as a single body, while for iC3b the TE domain was modelled as a rigid body and the cleaved CUB domain was modeled as connected dummy residues. For C3b all output models clustered closely, showing a slight detachment of the TE domain from the MG1 domain (Fig. S2E-F) consistent with prior results (38). For iC3b, 100 independent rigid body models were generated with two different initial positions of the TE domain. In the first initial model, the TE domain was positioned as in C3b, and in the other the TE domain was positioned next to the C345c domain. Independent of the initial position of the TE domain, the 10 models with the best fit to the experimental data clustered closely in terms of conformation. In these models, the cleaved CUB domain adopts an extended conformation with the TE domain positioned far from MG1 and the rest of the MG-ring (Fig. 3A). Apart from being detached from the MG-ring, the CUB-TE moiety has also rotated so that the TE domain is located towards the MG4 domain edge of C3c, whereas in crystal structures of C3b the TE domain forms contacts with the MG1 domain (Fig. 3A). In contrast, other models with a significantly worse fit to the data had the TE domain close to the C345c domain. A comparison of these two classes of rigid body models reveals that even though they are very different with respect to the location of the TE domain, their mass distribution is similar (Fig. 3C-D). In summary, the degraded CUB domain adopts an extended conformation, leading to the TE domain detaching significantly from the MG core to a preferred position rather than being randomly located relative to the C3c moiety.

**Figure 3.**
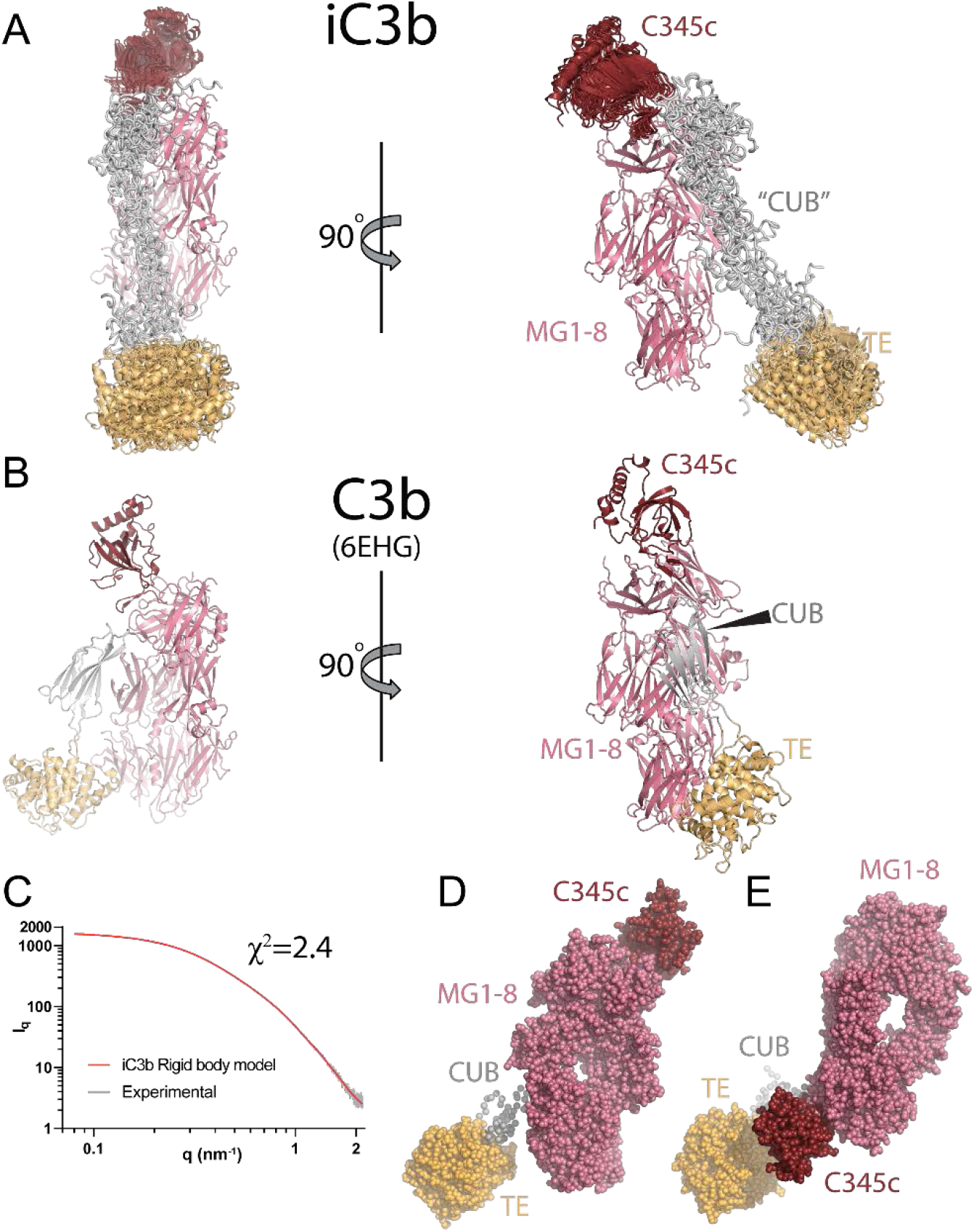
SAXS rigid body analysis of iC3b. **A.** The 10 rigid body models with the lowest χ^2^-value aligned on the MG1-6 domains. The models cluster closely together with their thioester domain displaced from the MG-ring, connected by an extended degraded CUB domain. C3b is presented in the same orientation for comparison. **B.** Experimental scattering curve (grey) compared with the curve calculated (red) from the iC3b rigid model shown in C. **C.** The iC3b model with the lowest χ^2^-value shown in a spheres representation. **D.** An alternative iC3b rigid body model where the thioester domain is in close proximity to the C345c domain. Notice how the mass distribution resembles that of the best fitting model in panel C.

### The iC3b-CR3 complex is compact

To characterize the solution structure of the iC3b:CR3 headpiece complex we collected synchrotron inline SEC-SAXS data. The forward scattering elution profile displayed two peaks - the first one corresponding to the complex, and the second one corresponding to excess iC3b (Fig. 4A). The R_g_ was stable throughout the first peak demonstrating that the CR3 remained saturated with iC3b during the SEC run in agreement with the 30 nM K_D_ observed by SPR. A Guinier analysis of the scattering curve did not indicate interparticle effects and suggested a R_g_ of 67 Å for the CR3-iC3b complex (Fig S3A). Based on calculation of the pair distribution function the D_max_ for the complex was ~260 Å (Fig. S3C). In support of a well-defined and compact CR3:iC3b complex, this is only slightly larger than the ~200 Å we observe for the iC3b and the CR3 headpiece monomers.

**Figure 4.**
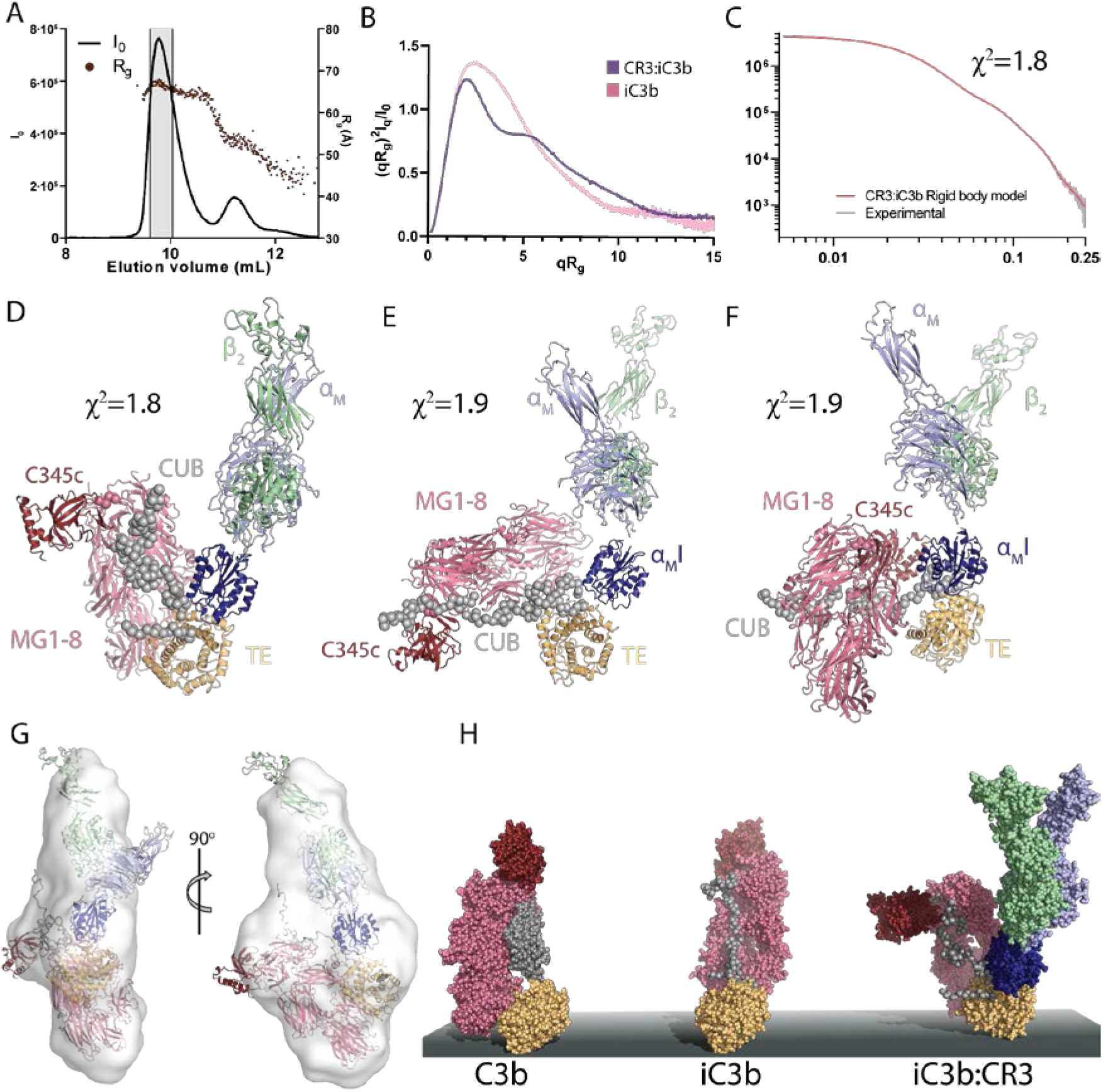
SEC-SAXS analysis of the iC3b:CR3 in complex. **A.** The forward scattering and R_g_ of each frame during the SEC-SAXS experiment of the CR3:iC3b complex plotted as a function of the elution volume. The two peaks contain the CR3:iC3b complex and excess free iC3b, respectively. The experimental scattering curve for the CR3:iC3b complex was obtained from the shaded area. **B.** The Guiner-normalized Kratky plot of the CR3:iC3b complex (purple) and iC3b alone (pink). **C.** Experimental scattering curve (grey) compared with the curve calculated (red) from the CR3:iC3b rigid model shown in panel D. **D-F.** Representative rigid body models of the three clusters of rigid body models. Upon CR3 recognition, the CUB remnants reorganizes, leading to a more compact iC3b conformation, but the exact position of C3c varies significantly between the three clusters. **G.** Average model of 32 ab initio models calculated from the CR3:iC3b SEC-SAXS data aligned to the rigid body model of the complex in panel D. **H.** Models of C3b, iC3b and the CR3:iC3b complex on an activator surface with coloring as in panels D-F. C3b is deposited on the surface and cleaved by FI into iC3b. The degraded CUB domain in iC3b extends releasing the C3c moiety of iC3b from the TE domain. Upon CR3 recognition, the CUB domain reorganizes substantially.

To obtain models of the iC3b:CR3 complex and to identify the parts of iC3b involved in the complex with CR3 apart from the TE domain, we performed rigid body refinement against the SAXS data. Within the complex, iC3b was modelled as for free iC3b except that the TE domain was fixed relative to the α_M_I domain of CR3, to maintain the interaction known from the crystal structure. The resulting rigid body was connected through two distance restrains between the α_M_I domain and the α_M_ β-propeller. This allows the α_M_I domain to rotate relative to the rest of the CR3 headpiece. The remaining fragment of the CR3 headpiece was modelled as a single rigid body in the open conformation. To improve the model fit to the data, the position of the C3c fragment in the best initial solutions was used as a new initial model, and the translation and rotation of the rigid bodies were sampled more finely. After two iterations, this strategy converged and resulted in the best model fitting the data with χ^2^=1.8 (Fig. 4C). The 15 best fitting models with 1.8<χ^2^<2.2 were classified into three tight clusters (Fig. 4D-F & Fig. S3D-F). In all the analyzed models, iC3b adopts a conformation markedly different from that of unbound iC3b. Relocation of the C3c moiety brings the iC3b TE domain close to the MG-ring (Fig. 4D-F) as opposed to unbound iC3b where the TE domain is located distantly from the MG-ring (Fig. 3A). However, our models do not allow us to define a unique position of the C3c fragment relative to the rest of the complex as rather different C3c orientations are observed in the three clusters of models (Fig. 4D-F & Fig. S3D-F).

The DAMMIF program was used to generate 40 ab initio models from the CR3:iC3b scattering curve. The models were subsequently clustered using DAMCLUST, which gave one major cluster containing 32 models. The models were aligned and averaged, which resulted in a flat and extended ab initio envelope, with similar dimensions to the rigid body models (Fig. 4G). Taken together, our SAXS data support a model where upon CR3 recognition, the C3c and TE domain moieties of iC3b are brought closer to one another because of recognition by CR3 of regions outside the TE domain. This results in a compact receptor-ligand complex which upon extrapolation to an opsonized cell recognized by CR3 predicts the C3c moiety of iC3b to be overall much closer to the activator rather than extended away from the surface as predicted for C3b (Fig. 4H).

## Discussion

Prior crystal structure and biophysical experiments of the α_M_I-C3d complex defined the core of the iC3b-CR3 interaction centered on the coordination of the divalent cation in the α_M_I MIDAS by an aspartate from the iC3b TE domain. It was further demonstrated that C3d and C3dg bound the α_M_ I-domain with an affinity resembling that of the iC3b-α_M_ I-domain interaction (30, 39). This C3d(g)-CR3 interaction appears to have physiological relevance since C3dg mediated erythrophagocytosis may occur in individuals suffering from paroxysmal nocturnal hemoglobinuria (39). However, other regions in CR3 beside the α_M_ I-domain must contribute to iC3b binding, since its deletion leaves residual iC3b affinity in CR3 (34). Both the α_M_ β-propeller and the β_2_ I-like domain have been implicated in interaction with iC3b (34, 36, 40, 41), and on the iC3b side, mutations in the iC3b Nt-α’ region associating with the MG7 domain weaken the iC3b-CR3 interaction (35). All these prior lines of evidence are consistent with a higher affinity of the CR3 headpiece for iC3b as compared to C3dg.

Prior studies have disagreed upon the affinity of CR3 for iC3b, and have been limited by a number of factors, including activation state of the receptor, purity of the protein preparations, and degree of oligomerization of the iC3b preparations (31, 32). We now quantitate the binding using highly pure and well-characterized proteins, and show that CR3 recognized iC3b with a K_D_ of 30 nM in Mg^2+^/Ca^2+^. This is 14-fold higher than the 515 nM observed for the CR3-C3d interaction in Mg^2+^/Ca^2+^, and is to our knowledge, the highest monovalent affinity measured between C3b, iC3b, C3dg and their five complement receptors. Although the difference in ΔG for these two dissociation constants only corresponds to 7 kJ/mol, due to multiple complexes formed between a phagocyte and an iC3b opsonized activator, a strong effect is predicted to result in vivo. Hence, if our findings translate to cell-bound CR3, the much more efficient phagocytosis of iC3b tagged objects compared to those tagged by C3dg can be rationalized. Importantly, it is only the combination of the CR3 headpiece and iC3b that results in this high affinity interaction, as both the CR3-C3d (this study) and the α_M_I-iC3b interactions (30) have K_D_ values of 500-600 nM. Thus, omission of the CUB and C3c moities of iC3b, or of the βI and β-propeller moieties of CR3, result in lower affinities, agreeing with our finding that these moieties come close to one another in iC3b-CR3 complexes.

Our rigid body models of the CR3:iC3b complex suggest that the C3c moiety of iC3b is in close proximity to CR3. A comparison with our models of unbound iC3b suggest that the TE domain is brought closer to C3c moiety upon CR3 binding. The heterogeneity in our SAXS models with respect to the location of the C3c moiety of iC3b relative to CR3 may reflect the in vivo situation. Alternatively, the limited information content present in the underlying SAXS data can give rise to rigid body models that from a structural point of view are quite different but fit the data equally well. The system is also challenging for SAXS rigid body refinement due to the difficulty of modeling the remnants of the CUB domain. Our refinement strategy was also conservative using only four rigid bodies to describe a 300 kDa complex with a D_max_ of 26 nm. Despite these limitations, our SAXS models are overall compatible with an ensemble of CR3-iC3b orientations seen in a recent nsEM study (2), except that those complexes showed more separation between the end of the C3c MG ring bearing the C345c domain and the CR3 β_I_-β-propeller interface. The authors suggested a second, three-dimensional contact formed between the integrin and iC3b that was less stable than the αMI-TE domain contact and was susceptible to disruption when iC3b-CR3 complexes adsorbed to the grid and become largely planar. These results are consistent with lack of co-planarity of the MG ring of C3c and the integrin headpiece in our rigid body models. We have also been unable to observe compact iC3b-CR3 complex particles using nsEM grids despite extensive efforts and use of gradient fixation (42).

The structural arrangement of iC3b has been somewhat controversial, with prior SAXS and nsEM studies disagreeing both within and between methods (2–4, 43, 44). In most EM-based studies, the TE domain was observed to be flexibly attached to the C3c moiety (2–4, 44). However, in one EM study the TE domain was found stably associated with the C345c domain (43). This study also presented SAXS data with significantly lower values of D_max_ and R_g_ than those observed by us and others (3, 43). Our data are more in line with other EM-based models of iC3b, where the TE domain is loosely associated with the C3c moiety through an extended CUB domain (2–4, 44).

Inhibition of specific CR3-ligand interactions has been investigated for decades, but is complicated by the plethora of structurally diverse CR3 ligands reported. Multiple CR3 function blocking antibodies are known, e.g. (45–47) and small molecules known as leukadherins binding CR3 and suppressing outside-in signaling upon ligand binding reduce inflammation and suppress tumor growth in animal models of cancer (48, 49). Recent developments in neurobiology are likely to fuel the interest for an iC3b-specific CR3 inhibitor. During development, activation of the classical pathway of complement on weakly signaling synapses leads to iC3b deposition and recognition by CR3-expressing microglia, which phagocytize the iC3b opsonized synapses (8, 9, 50). Very recently, microglia CR3 was shown to support complement-dependent synapse elimination by microglia as a mechanism underlying the forgetting of remote memories (51). However, the same pathway that ensures correct development and removal of remote memories by pruning excess synapses, is linked to Alzheimer’s disease (13), frontotemporal dementia (FTD) (50) and spinal muscular atrophy (52). Our demonstration of a stable and compact complex between iC3b and the CR3 headpiece with a dissociation constant in the low nanomolar range offers hope for the development of molecules capable of specifically interfering with the iC3b:CR3 interaction while preserving the ability of CR3 to recognize its many other non-complement ligands.

## Materials and methods

### Generation of a stable cell line expressing the CR3 headpiece fragment

The coding sequence of the human CR3 α_M_-chain residues 17-773 containing the glycan knockout mutations N225R/N680R and β_2_-chain residues 23-504 were cloned into the pIRES2-EGFP based in-house vectors ET10c and ET10b respectively. The ET10c vector contains a Human Rhinovirus (HRV) 3C protease recognition site, an acid coiled-coil region, a StrepII-tag and a His_6_-tag on the 3’, directly in-frame with the cloning site. The open reading frame was subsequently subcloned into pcDNA3.1(+). The ET10b vector contains an HRV 3C protease recognition site, a basic coiled-coil region, and a His_6_-tag directly in-frame with the cloning site. The CR3 α_M_- and β_2_-chain were co-transfected into human embryonic kidney (HEK) 293S GnTi^−^ cells (ATCC). The selection antibiotics Hygromycin B and G418 at 200 µg/mL and 1g/mL, respectively, were added to the cultures 48 hours post transfection. After selection the cells were assessed for GFP expression using fluorescence-activated cell sorting, and the top 5 % expressing clones were seeded in a 96 well cell culture plate. A final selection step was performed on the cell supernatants using sandwich ELISA by capturing the CR3 headpiece by use of an anti-CR3 α_M_-chain antibody (CBRM 1/2), and detected using a biotinylated anti-CR3 β_2_-chain antibody (IB4).

### Expression and purification of CR3 headpiece

The CR3 headpiece stably transfected HEK293S cells were kept as adhesion cell culture growing in Dulbecco’s Modification of Eagle’s Medium (DMEM) GlutaMAX (Gibco) supplemented with 10 % (v/v) fetal bovine serum (FBS), 20 mM HEPES pH 7.5, 1 % Penicillin-Streptomycin (Gibco), 200 µg/mL Hygromycin B (Sigma-Aldrich) and 200 µg/mL G418 (Sigma-Aldrich). Before large scale purify cation, the cells were adapted to serum-free medium. The cell supernatant was harvested by centrifugation and subsequently filtered through 0.2 µm filters. The cleared cell supernatant was supplemented with 50 mM TRIS pH 8, 500 mM NaCl, 5 mM MgCl_2_ and 1 mM CaCl_2_ and applied to a 5 mL HisTrap Excel (GE Healthcare). Afterwards the column was washed with 40 mL of 20 mM TRIS pH 8, 1.5 M NaCl, 5 mM MgCl_2_, 1 mM CaCl_2_ and the protein was eluted in 20 mL of 20 mM TRIS pH 8, 150 mM NaCl, 5 mM MgCl_2_, 1 mM CaCl_2_, 400 mM imidazole. The elution was applied to a 1 mL StrepTactin column (GE Healthcare) equilibrated in 20 mM HEPES pH 7.5, 150 mM NaCl, 5 mM MgCl_2_, 1 mM CaCl_2_. The column was washed in 20 mM HEPES pH 7.5, 150 mM NaCl, 5 mM MgCl_2_, 1 mM CaCl_2_ and the protein was subsequently eluted in 20 mM HEPES pH 7.5, 150 mM NaCl, 5 mM MgCl_2_, 1 mM CaCl_2_, 2.5 mM D-desthiobiotin. 3C rhinovirus protease was added in a 1:10 mass ratio to CR3 and the reaction was allowed to proceed at 4°C overnight. A final polishing step was performed by size exclusion chromatography (SEC) on a 24 mL Superdex 200 increase (GE Healthcare) equilibrated in 20 mM HEPES pH 7.5, 150 mM NaCl, 5 mM MgCl_2_ and 1 mM CaCl_2_.

### Surface plasmon resonance assays

Human C3d was expressed and purified as described in (30). C3b and iC3b was generated and purified as described in (37). The experiments were performed on a Biacore T200 instrument with a running buffer containing 20 mM HEPES pH 7.5, 150 mM NaCl, 5 mM MgCl_2_, 1 mM CaCl_2_ unless otherwise stated. Streptavidin was immobilized on a CMD500M chip (XanTec Bioanalytics) to 200 response units. C3d, or iC3b biotinylated on the thioester cysteine was injected on the chip until the surface was saturated. For the kinetics experiments using iC3b, the CR3 headpiece was injected in a concentration series ranging from 0.3215 nM to 100 nM, whereas for C3d, the CR3 headpiece was injected in a concentration series ranging from 3.25 nM to 2000 nM. The surface was regenerated by using a buffer containing 50 mM EDTA, 1 M NaCl, 100 mM HEPES pH 7.5. The data were analyzed using a 1:1 binding model, and the reported on- and off-rates are averages of three independent experiments. The kinetic experiment with iC3b on the surface was repeated three times in the buffer containing 20 mM HEPES pH 7.5, 150 mM NaCl, 1 mM MnCl_2_, 0.2 mM CaCl_2_ as well. The competition assays were performed on the iC3b surface where 20 mM of CR3 headpiece was injected either alone, or pre-incubated on ice for 1 hour with 10, 20, 50, 100, 200, or 1000 nM of iC3b, C3d or C3b respectively. All experiments were performed in triplicates.

### Analytical SEC analysis

For analyzing the effect of different divalent cations on the oligomeric state of CR3, 50 µL of CR3 headpiece at 2 µg/µL was diluted four-fold in either 20 mM HEPES pH 7.5, 150 mM NaCl, 1 mM MnCl_2_, 0.2 mM CaCl_2_; 20 mM HEPES pH 7.5, 150 mM NaCl, 5 mM MgCl_2_, 1 mM CaCl_2_; or 20 mM HEPES pH 7.5, 150 mM NaCl, 5 mM NiCl_2_, 1 mM CaCl_2_. The protein was incubated for 1 hour at room temperature before being injected on a 24 mL Superdex 200 increase equilibrated in the respective protein dilution buffer. For analyzing the complex formation between CR3 and iC3b, 15 µg of iC3b was mixed with 1.1 fold molar excess of CR3. The sample was injected on a 2.4 mL Superdex 200 increase equilibrated in 20 mM HEPES pH 7.5, 150 mM NaCl, 5 mM MgCl_2_, 1 mM CaCl_2_. Control experiments injecting either CR3 or iC3b in the same amount was also performed.

### SAXS data collection and analysis

In-line SEC-SAXS data for the CR3 headpiece in complex with iC3b were collected at the P12 beamline at PETRA III, Hamburg, Germany (53). Scattering was recorded from the elution of a 24 mL Superdex 200 increase equilibrated in 20 mM HEPES pH 7.5, 150 mM NaCl, 5 mM MgCl_2_, 1 mM CaCl_2_ with a flow rate of 0.25 mL/min. The CR3 headpiece was mixed with 1.2 fold molar excess of iC3b and subsequently injected on the SEC column. Each frame during the SEC-SAXS run covers a 0.955 s exposure performed every second. The sample-to-detector distance was 3.0 m covering 0.02 < *q* < 4.8 nm^−1^ (q=4π·sinθ·λ^−1^, where 2θ is the scattering angle). Normalization and radial averaging was performed at the beamline using the automated pipeline (53, 54). Buffer subtraction was performed using CHROMIXS (55). Due to the high level of averaging performed during SEC-SAXS data processing, detector imperfections most likely led to the error estimates being underestimated at low q, as revealed an indirect Fourier transform of the data. The error estimates were therefore rescaled by Err=Err_original_ ·(1+2.5·exp(−7q^2^)). Forty ab initio models were generated using dammif in slow-mode, and the resulting models were clustered using damclust (56). This lead to six clusters with only one major cluster containing 32 of the 40 models. The models of each cluster was subsequently averaged and filtered yielding the final models of each cluster. SAXS measurements of CR3 headpiece, C3b and iC3b in batch mode, were likewise collected at the P12 beamline. The data were collected in a temperature-controlled capillary at 20°C using a PILATUS 2M pixel detector (DECTRIS) with λ = 1.240 Å. The sample-to-detector distance was 3.0 m covering 0.02 < *q* < 4.8 nm^−1^. Samples of CR3 were prepared at 0.7, 1.3, 1.7 and 2.9 mg/mL, and samples of C3b, and iC3b were prepared at 7.3, and 13.0 mg/mL respectively, where after data was collected with twenty exposures of 45 ms. Radial averaging, buffer subtraction and concentration scaling was performed by the automated beamline pipeline (57) and the pair distribution function was calculated by indirect Fourier transformation using GNOM (58).

### SAXS rigid body modelling

Rigid body modelling using CORAL (56), was performed for C3b, iC3b and the CR3:iC3b complex against their corresponding SAXS curves. Input domains for C3b and iC3b was extracted from RCSB entry 6EHG (37), after aligning the principal inertial vectors of the structure with the Cartesian axes using alpraxin (59). For C3b the MG1-6, MG7, C345c and the CUB-TE fragment were extracted as separate files. The rigid bodies were connected using LINK statements to model the missing linker regions. The MG1-7 and MG8 rigid bodies were fixed relative to each other such that the MG ring was kept intact, and a distance restraint of 6 Å was applied between Cys873 in the MG8 domain and Cys1513 in the C345c domain to maintain their disulphide bridge. Twenty rigid body models were generated with CORAL, which are all shown in fig. S2E. For iC3b a similar scheme was used, except that the CUB domain was modelled using dummy residues to represent all three flexible stretches of the degraded domain. This was done by (1) introducing a LINK statement connecting the MG7 C-terminal residue to the N-terminal residue of the TE domain. (2) Introducing a CTER statement at the C-terminal residue of the TE domain. (3) Introducing a NTER statement at the N-terminal residue of the MG8 domain. One hundred coral runs were performed, half using an input model where the TE domain was placed in a C3b like conformation and the other half where the TE domain was placed beside the C345c domain. No systematic differences could be observed between the models coming from either input models. The ten models with the best fit to the data are shown in Fig. 3A. For the CR3:iC3b complex iC3b was modelled as for free iC3b, except that the TE domain was fixed relative to the α_M_I domain of CR3, to keep the interface known from the crystal structure of the complex (RCSB entry 4M76 (60)) intact. The N- and C-terminal residues of the CR3 α_M_I was restrained with two distance restraints of either 7 or 9 Å to the α_M_ β-propeller since with distance restraints of 5 Å very few models obtained χ^2^ < 3. The rest of the CR3 headpiece was kept as a single rigid body, using a homology model based on α_X_β_2_ (4NEH), modelled in the open conformation using the β-chain from the crystal structure of α_IIb_ β_3_ (RCSB entry 2VDR (61)). Three consecutive rounds of rigid body modelling were performed, in rounds 2 and 3 starting from the best fitting model of the prior round. To sample the rigid body movement more finely, the CORAL parameters “angular step” and “spatial step” were decreased from the default 20° and 5 Å to 10° and 2.5 Å respectively during optimization. No systematic differences could be observed between the models coming from different input models and subject to 7 or 9 Å distance restraints. Hence, a total of 135 output models were analyzed and the best 15 models having 1.8<χ^2^<2.2 were analyzed.

## Acknowledgements and funding sources

We thank the staff at the P12 beamline at PETRAIII for help during data collection, and Lise Arleth and Jan Skov Pedersen for discussions on correction of SAXS data. The authors would like to acknowledge Christine Schar for assistance with SPR and Karen Margrethe Nielsen for technical support. This work was supported by the Lundbeck Foundation (BRAINSTRUC, grant no. R155-2015-2666) and the Danish Foundation for Independent Research (grant no 4181-00137).

