## Supporting information for "Complement receptor 3 forms a compact high affinity complex with iC3b"

Supporting information figures S1-S3

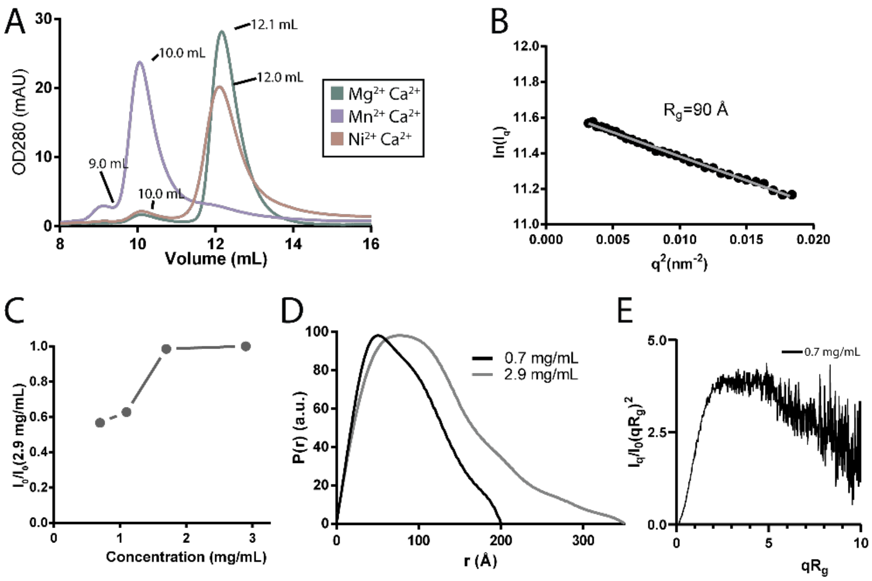


**Supporting information figure 1. CR3 dimerizes in a concentration dependent manner. A.** Comparison of analytical SEC runs with the CR3 headpiece fragment in $MgCl_2$ (turquoise), $MnCl_2$ (purple), and $NiCl_2$ (brown), revealing the cation dependence of CR3 headpiece oligomerization. **B.** Guinier analysis of the CR3 headpiece SAXS data at 2.9 mg/mL. **C.** The forward scattering of CR3 is plotted as a function of concentration, indicating that CR3 undergoes dimerization at higher concentrations. **D.** Pair distribution functions of CR3 headpiece at either 0.7 mg/mL (black) or 2.9 mg/mL (grey) showing that $D_{max}$ increases from ~200 to 350 Å at higher concentrations. **E.** The Guinier normalized Kratky plot of the CR3 headpiece at 2.9 mg/mL showing that CR3 is an ordered protein with some flexibility.

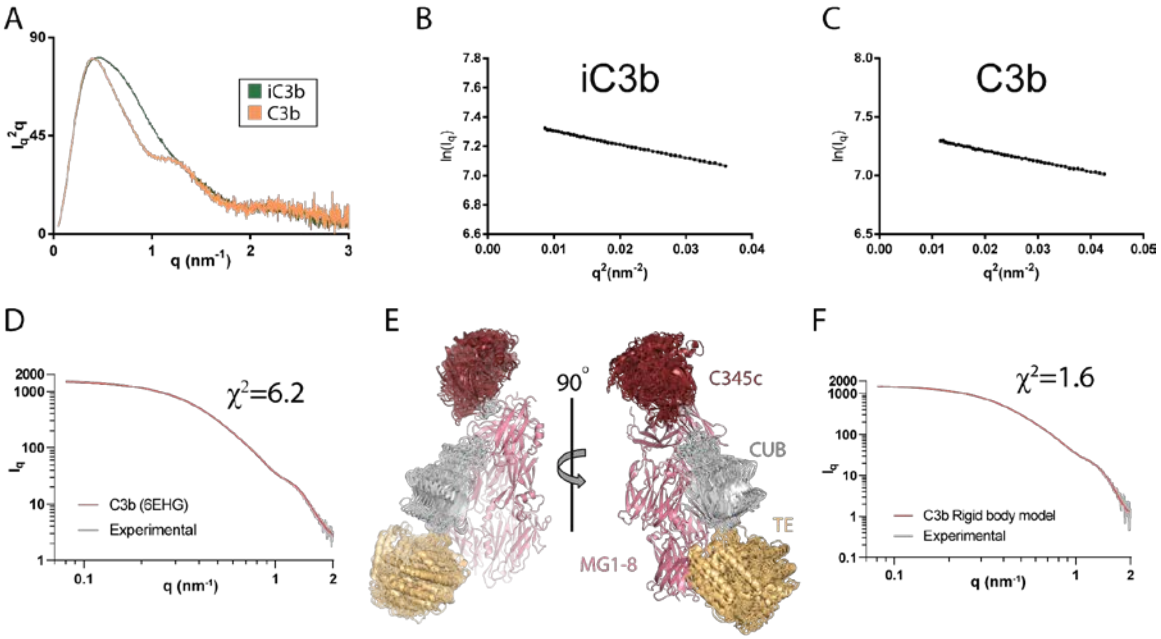


**Supporting information figure 2. SAXS analysis of iC3b and C3b. A.** Comparison of the scattering curve of iC3b and C3b plotted as a Kratky plot with $I_q \cdot q^2$ plotted against q, showing that iC3b adopts a more extended conformation than C3b. **B-C**. Guinier plots of iC3b and C3b with the experimental points plotted in black and a linear regression fit in grey. **D.** The experimental scattering of C3b compared to the calculated scattering of C3b (RCSB entry 6EHG) calculated with crysol. This demonstrates that the crystal structure is close to the solution conformation of C3b. **E.** Twenty rigid body models of C3b calculated with coral, aligned on the MG1-8 domains. All models cluster closely with the TE domain detached slightly from the MG-core as compared to the crystal structure. **F.** The experimental scattering of C3b compared to the calculated scattering of one of the C3b rigid body models presented in panel E.

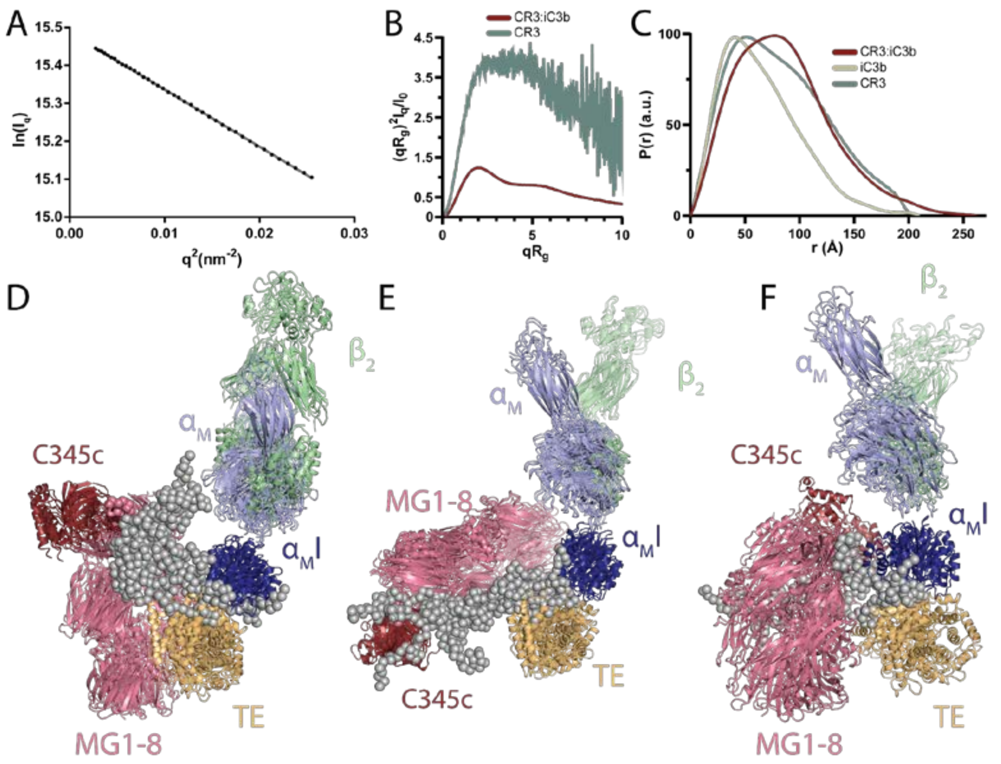


**Supporting information figure 3. SAXS analysis of the CR3:iC3b complex. A.** Guinier plot of the scattering curve for the CR3:iC3b complex. **B.** Comparison of the Guinier normalized Kratky plots of the CR3:iC3b complex (red) and CR3 headpiece (turquoise). **C**. The pair distribution function of the CR3:iC3b complex suggesting a $D_{max}$ of 26 nm for the CR3:iC3b complex. The complex is compared to iC3b (sand) and CR3 (blue) which both have $D_{max}$ values of ~20 nm. **D-F.** All models from the three clusters of CR3:iC3b rigid body models.

| Receptor | Ligand | Reference | $k_{on}$ ($M^{-1}$ $s^{-1}$) | $k_{off}$ ($s^{-1}$) | $K_D$ (nM) |
|---|---|---|---|---|---|
| CR3 HP | iC3b ($Mg^{2+}$) | This study | $1.22 \cdot 10^6 \pm 0.44 \cdot 10^6$ | $3.60 \cdot 10^{-2} \pm 0.035 \cdot 10^{-2}$ | 29.6 ± 10 |
| CR3 HP | iC3b ($Mn^{2+}$) | This study | $1.95 \cdot 10^5 \pm 0.029 \cdot 10^5$ | $1.21 \cdot 10^{-3} \pm 0.016 \cdot 10^{-3}$ | 6.17 ± 0.11 |
| CR3 HP | C3d ($Mg^{2+}$) | This study | | | 515 ± 33.8 |
| $\alpha_M$I | iC3b($Mg^{2+}$) | (1) | | | 600 |
| $\alpha_M$I | C3d ($Mg^{2+}$) | (1) | | | 450 |
| CR3 | iC3b | (2) | | | 210 |
| CR3 | iC3b ($Mg^{2+}$) | (3) | | | 12.5 |

**Supporting information table 1. Comparison of CR3 headpiece interactions with various C3 fragments to prior research.** For our SPR data, the on- and off-rates are shown as average values of three independent experiments ± the standard deviation. For C3d steady-state analysis was performed to calculate the dissociation constant, which was obtained by non-linear regression against the average equilibrium binding response of three independent experiments, and the $K_D$ is reported ± the standard deviation of the fit.
